# Specificities and commonalities of the Planctomycetes plasmidome

**DOI:** 10.1101/2024.01.22.576637

**Authors:** María del Mar Quiñonero Coronel, Damien Paul Devos, M. Pilar Garcillán-Barcia

## Abstract

Although plasmids play a crucial role in antibiotic resistance and contemporary biotechnology, our comprehension of their natural ecological dynamics remains restricted to a few bacterial groups. Planctomycetes is a bacterial phylum with unusual molecular and cellular biology for which little is known about its plasmidome. This study provides the first comprehensive description of the diversity of the endogenous plasmids found in Planctomycetes, which may be used as starting points to generate different genetic tools for research on this phylum. Plasmids from Planctomycetes encode a wide variety of biological functions, although for a large portion of the encoded genes a function cannot be assigned, which is usual in Planctomycetes. They seemed to have largely coevolved with the genome of their hosts, with which they share many homologs. We also detected recent transfer events of insertion sequences between co-habiting chromosomes and plasmids. 60% of the plasmid genes are distantly related to chromosomally-encoded genes and 40% have homologs in plasmids from other bacterial groups, while 36% of the proteins composing the planctomycetal plasmidome are exclusive. Most planctomycetal plasmids encode a replication initiation protein of the RPA family in the proximity of a putative iteron-containing replication origin, as well as active type I partition systems. One conjugative and three mobilizable plasmids were identified, suggesting horizontal gene transfer via conjugation in this phylum.

## INTRODUCTION

Most of our current knowledge on the bacterial world is largely based on model organisms from a restricted subset of phyla, mostly Proteobacteria and Firmicutes. Within this paradigm, notable exceptions have emerged, showcasing a rich tapestry of evolutionary adaptations. The phylum Planctomycetota (formerly Planctomycetes) stands out as one of the most enigmatic branches within the bacterial evolutionary tree. Planctomycetes exhibit a distinctive cellular biology replete with attributes seldom encountered in the bacterial domain, thereby piquing scientific interest due to their potential relevance to the eukaryogenic process (Forterre, 2011; Devos, 2021). Intriguinly, they manifest an atypical cellular configuration characterized by extensive invaginations of the cytoplasmic membrane, forming a complex endomembrane system (Acehan et al., 2013; Devos, 2014; Forterre, 2011; Santarella-Mellwig et al., 2013), previously mistaken as cell compartmentalization (Lindsay et al., 2001). Some of the cells display a phenotype similar to the typically eukaryotic characters of endo– and phago-cytosis (Lonhienne et al., 2010; Shiratori et al., 2019). Furthermore, they encode homologs to several eukaryotic proteins (Jenkins et al., 2002; Pearson et al., 2003; Santarella-Mellwig et al., 2010; Makarova and Koonin, 2010; Arcas et al., 2013; Shiratori et al., 2019; Santana-Molina et al., 2020). But beyond their distinctive cellular attributes, Planctomycetes are recognized as pivotal contributors to global carbon and nitrogen cycles, owing to their metabolic proficiency in the degradation of complex carbon substrates (Jeske et al., 2013; Wiegand et al., 2018), and their unique ability to combine ammonium and nitrite or nitrate to form nitrogen gas (Kuenen, 2008; Strous et al., 1999). Moreover, their genomic repertoire encompasses genes associated with secondary metabolite pathways, which are implicated in the production of bioactive compounds, including antimicrobial and algicidal agents (Jeske et al., 2013, 2016; Graça et al., 2016).

Metagenomic data indicated the ubiquity of Planctomycetes in diverse ecosystems (Wiegand et al., 2018), encompassing soil (Stackebrandt et al., 1993; Wang et al., 2002; Buckley et al., 2006; Ivanova et al., 2016;), plant rhizosphere (Lei et al., 2023; Ravinath et al., 2022), seawater (Schlesner, 1994; Wiegand et al., 2020), freshwater (Schlesner, 1994; Wang et al., 2002; Dedysh et al., 2020), chlorinated water (Aghnatios and Drancourt, 2015), the human gastrointestinal tract (Cayrou et al., 2013), and very frequently in association with several algae (Bengtsson and Øvreås, 2010; Lage and Bondoso, 2011, 2012; Wiegand et al., 2020). However, the cultivation of Planctomycetes in pure axenic cultures has proven to be challenging, primarily attributed to their recalcitrance to grow on synthetic media and their inherently slow growth kinetics (Kaboré et al., 2020). Only a minute fraction, approximately 0.6%, of known planctomycetal operational taxonomic units are successfully isolated (Wiegand et al., 2018). Besides, only a few examples of genetic modifications have been carried out in this phylum, mostly in *Planctopirus limnophila* (formerly known as *Planctomyces limnophilus*), and to a lesser extend in *Gemmata obscuriglobus*, *Gimesia maris*, and *Blastopirellula marina* (Rivas-Marín et al., 2016). Foreign DNA incorporation has been accomplished through electroporation (Jogler et al., 2011) and conjugation (Rivas-Marín et al., 2016), while the generation of mutants was carried out by homologous recombination (Erbilgin et al., 2014; Rivas-Marín et al., 2016) or transposon mutagenesis, employing the EZ-Tn*5* system (Jogler et al., 2011; Schreier et al., 2012; Rivas-Marín et al., 2023).

In 2020, a concerted effort was made to selectively isolate axenic cultures from all major planctomycetal clades, subsequently culminating in the sequencing of their entire genomes (Wiegand et al., 2020). The current accessibility of this genome dataset facilitates a thorough genomic analysis of this phylum. Notably, one of the significant research gaps in this domain pertains to the comprehensive exploration of plasmids within the Planctomycetes on a global scale. Plasmids are nearly ubiquitous in bacteria, and Planctomycetes are no exception to this rule (Guo et al., 2014; Ivanova et al., 2017; Jogler et al., 2020; Kulichevskaya et al., 2020). These genetic elements play a pivotal role in mediating genetic exchange in bacteria (Halary et al., 2010). Besides, the absence of replicative vectors remains a notable limitation to genetically manipulate Planctomycetes. Plasmids are a natural source to construct replicative vectors, which, in turn, are essential to enhancing the construction of deletion mutants and the development of expression systems. In the present study, we delve into the diversity within the planctomycetal plasmidome, elucidating its unique gene families, as well as those that are shared with plasmids originating from taxonomic groups outside this phylum or the planctomycetal host genomes. Moreover, we scrutinize the main characteristics of the planctomycetal plasmid modules, serving as an initial step for the prospective design of replicative vectors.

## RESULTS AND DISCUSSION

### Planctomycetal plasmids coevolve with their hosts

The scarcity of genome sequence data for many bacterial phyla is one of the main impairments to assess the contribution of plasmids, a fundamental part of the accessory genome, to their host adaptation, fitness and survival. The current planctomycetal plasmidome encompasses 21 plasmids distributed in 10 isolates (Supplementary Table S1). As expected because of its larger representation in the dataset, class *Planctomycetia* gathered most of the plasmids (20), mainly in hosts of the *Isosphaeraceae* family (order *Isosphaerales*) (Figure 1). Notably, all members of this taxonomic family contained plasmids, varying in number from one to five. The size of plasmids associated with Planctomycetes varies, ranging from 12.5 to 278 kb with a median size of 81.1 kb, following a unimodal distribution (Figure 2A).

**Figure 1.**
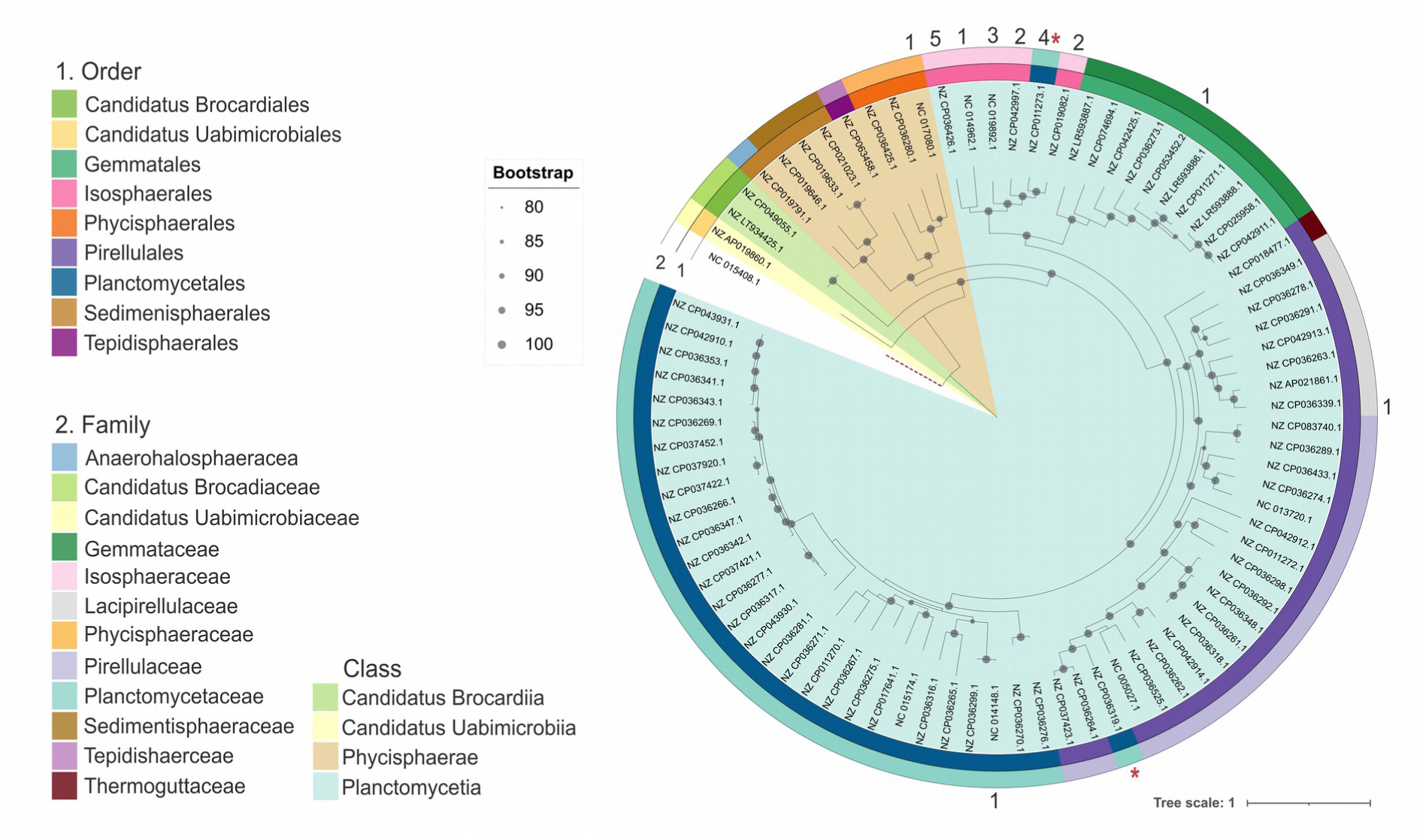
Plasmid distribution on the *Planctomycetota* phylogeny. ML phylogeny based on universal bacterial markers retrieved with PhyloPhlAn from the 83 planctomycetal chromosomes. The outgroup (*Chlamydia pecorum* E58, NC_015408.1) is indicated by a discontinuous red line. Grey circles are placed on the branches supported by ultrafast bootstrap values >=80%. The taxonomic metadata was retrieved from GenBank. The background color indicates the *Planctomycetota* classes, while taxonomic orders and families are shown in the inner and outer colored rings, according to the legend. The number of plasmids contained in the corresponding host is indicated. Incongruencies in the phylogenetic position of two isolates regarding the taxonomic metadata retrieved from GenBank are indicated by asterisks. These differences were already reported. *Crateriforma conspicua* strain Mal65 (NZ_CP36319.1) was described as a member of the *Planctomycetaceae* family (order *Planctomycetales*) by 16S rRNA phylogeny (Peeters et al., 2020) and as a member of the *Pirellulaceae* family (order *Pirellulales*) by a multilocus-sequence analysis-based tree (Vitorino and Lage, 2022), while *Planctomyces* sp. SH-PL62 (NZ_CP011273.1) was acknowledged as a member of the *Isosphaeraceae* family (order *Isosphaerales,* formerly *Planctomycetales*) by 16S rRNA phylogeny (Ivanova et al., 2017).

**Figure 2.**
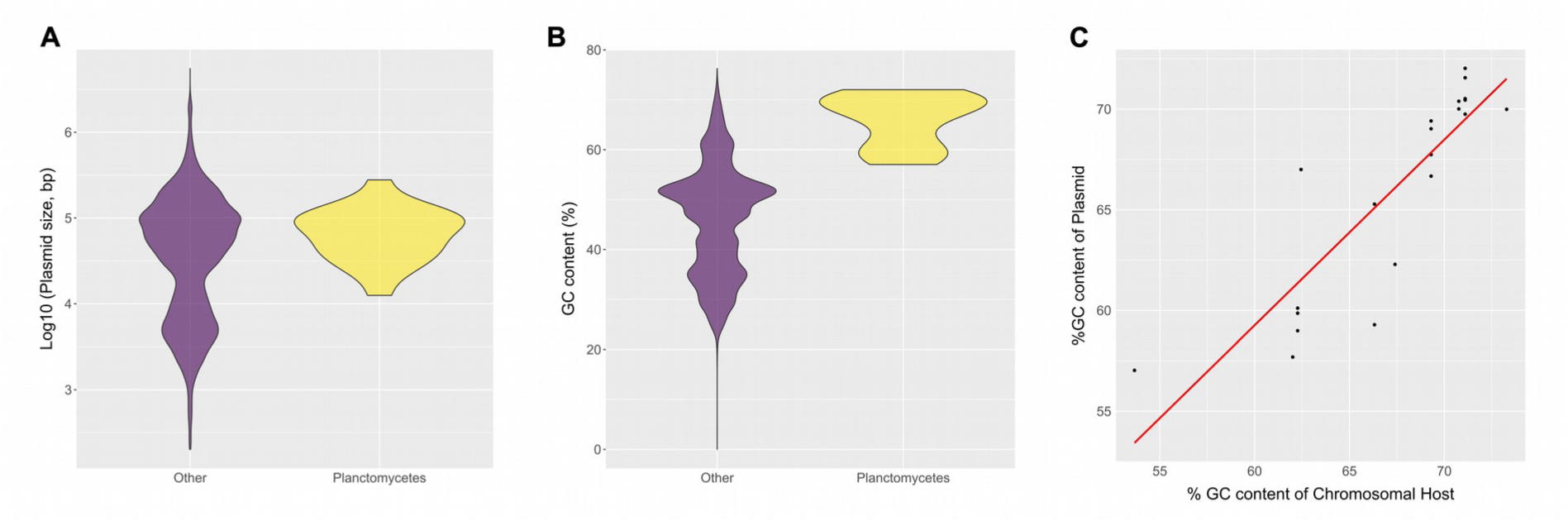
Comparison of planctomycetal plasmids with those from other bacterial phyla and their host genomes. Distribution of the planctomycetal plasmids (yellow) versus plasmids from other phyla (purple) by A) size and B) GC content. C) GC correlation between planctomycetal plasmids and their host genomes. The red line indicates the linear regression model of the data.

The GC content of planctomycetal plasmids is relatively higher compared to plasmids found in most other phyla, with values ranging from 57% to 72% and a median GC content of 67.7% (Figure 2B). Furthermore, there is a significant correlation between the GC content of planctomycetal plasmids and that of their respective host chromosomes, as evidenced by a high Pearson’s correlation coefficient of 0.86 (p-value = 4.72e-07) (Figure 2C). This pattern aligns with the general trend observed in plasmid-host relationships in other phyla, as described previously (Almpanis et al., 2018). Since sequences recently introduced into a bacterial genome often bear unusual sequence characteristics and gradually tend to adopt the average nucleotide composition of the host genome (Lawrence and Ochman, 1998), the GC similarity between plasmids and their hosts is an indicator of long co-evolution between them. As also previously observed in plasmids originated from other phyla (Almpanis et al., 2018; Nishida, 2012), the planctomycetal plasmids are richer in AT than their hosts, on average 1.5% higher. A higher energy cost and limited availability of G and C over A and T/U has been suggested as a basis for these differences (Rocha and Danchin, 2002).

### Biological functions encoded in the Planctomycetes plasmidome are very diverse

To explore the functions associated with plasmids hosted in Planctomycetes, the plasmid-encoded proteins (1,375) were mapped to the Protein Families (Pfam) and Cluster of Orthologous Group (COG) databases. 692 of them were assigned to 456 Pfam families (Supplementary Figure S1A) and 653 proteins to 20 out of the 23 different COG categories (Supplementary Figures S1B and S2), including 155 proteins assigned to the poorly characterized COG category S of ‘function unknown’, and other 53 proteins assigned to COGs without any category (Supplementary Table S2). Proteins assigned to Pfam families per plasmid ranged from 17 to 79% and those assigned to COGs from 12 to 96% (Supplementary Figure S1). Taking both annotation methods together, 57% of the plasmid proteome could be assigned to a protein family (Pfam and/or COG) with a known function. Proteins involved in replication, recombination and repair (129); membrane biogenesis (76); transcription (63); signal transduction (46) and inorganic ion transport and metabolism (44) were the most frequent COGs detected, and most of them were found more than once per plasmid (Supplementary Table S2).

Planctomycetes have been reported to specialize in sugar and secondary metabolism, with a notable emphasis on the degradation of high molecular weight sugars. This specialization is attributed to the presence of gene clusters encoding carbohydrate-active enzymes (CAZymes) (Andrade et al., 2017). Remarkably, it has been conjectured that the presence of these gene clusters may be a contributing factor to the relatively larger genomes observed within this taxonomic phylum. Notably, the COG analysis identified 20 proteins in 8 plasmids as putative CAZymes: 4 glycosyl hydrolases (GH) and 16 glycosyl transferases (GT) belonging to 4 and 6 CAZy families, respectively (Supplementary Table S2). These proteins are included in COG categories related to carbohydrate transport and metabolism (4 proteins belonging to COG category G); cell wall, membrane and/or envelope biogenesis (15 proteins to COG M), and energy production and conversion coupled with carbohydrate transport and metabolism (1 protein to COG G). The most abundant CAZy families detected were GT26 (n=5), GT4 (n=4), GT2,GT4 (n=3), and GT2 (n=2). A higher proportion of CAZymes encoded in plasmids are detected in hosts exhibiting a lower number of chromosomally-encoded CAZymes, which suggests that plasmids are likely complementing these metabolic pathways (Supplementary Table S3). An exceptional illustration can be found in the case of the host *Tautonia plasticadhaerens* EIP (NZ_CP036426.1), which encompasses 7 CAZymes within its chromosome. Additionally, it accommodates five plasmids, three of which, namely pEIP_2 (NZ_CP036428.1), pEIP_4 (NZ_CP036430.1), and pEIP_5 (NZ_CP036428.1), were identified to contain 1, 7, and 3 CAZymes, respectively.

### The currently available planctomycetal plasmidome is very diverse and its members share only a few homologs

Bacterial plasmids have been recently classified in Plasmid Taxonomic Units (PTUs), based on their average nucleotide identity, ANI_L50_ (Garcillán-Barcia et al., 2023; Redondo-Salvo et al., 2020). To analyze the similarity of Planctomycetes plasmids, we first calculated the pairwise ANI_L50_ values between them. Notably, despite prior observations of homologous regions and synteny conservation in certain plasmids of the *Isosphaeraceae* planctomycetes (Ivanova et al., 2017), all pairwise ANI_L50_ values equaled 0, indicating a high level of diversity in the plasmid backbones.

Subsequently, we undertook a comprehensive analysis of the entire open reading frame dataset (ORFeome) derived from the 21 planctomycetal plasmids, a total of 1,375 plasmid-encoded proteins, employing a clustering approach at 30% identity and 60% coverage, with the aim of identifying remotely related homologs. This analysis yielded 1,053 distinct homologous protein clusters (HPCs). To visually represent the relationships between plasmid genomes and HPCs, we constructed a bipartite network, as illustrated in Figure 3. In this network, connections between the two types of nodes (plasmid genomes and HPCs) were established when a member of a given protein cluster was present in a particular genome. As a result, plasmids sharing a greater number of HPCs tended to cluster together within the network. It is noteworthy that the resulting network displayed a sparse distribution, with only 135 protein clusters being shared by at least two plasmids, depicted as black nodes in Figure 3A. Conversely, 918 HPCs remained as singletons, each linked solely to a specific plasmid.

**Figure 3.**
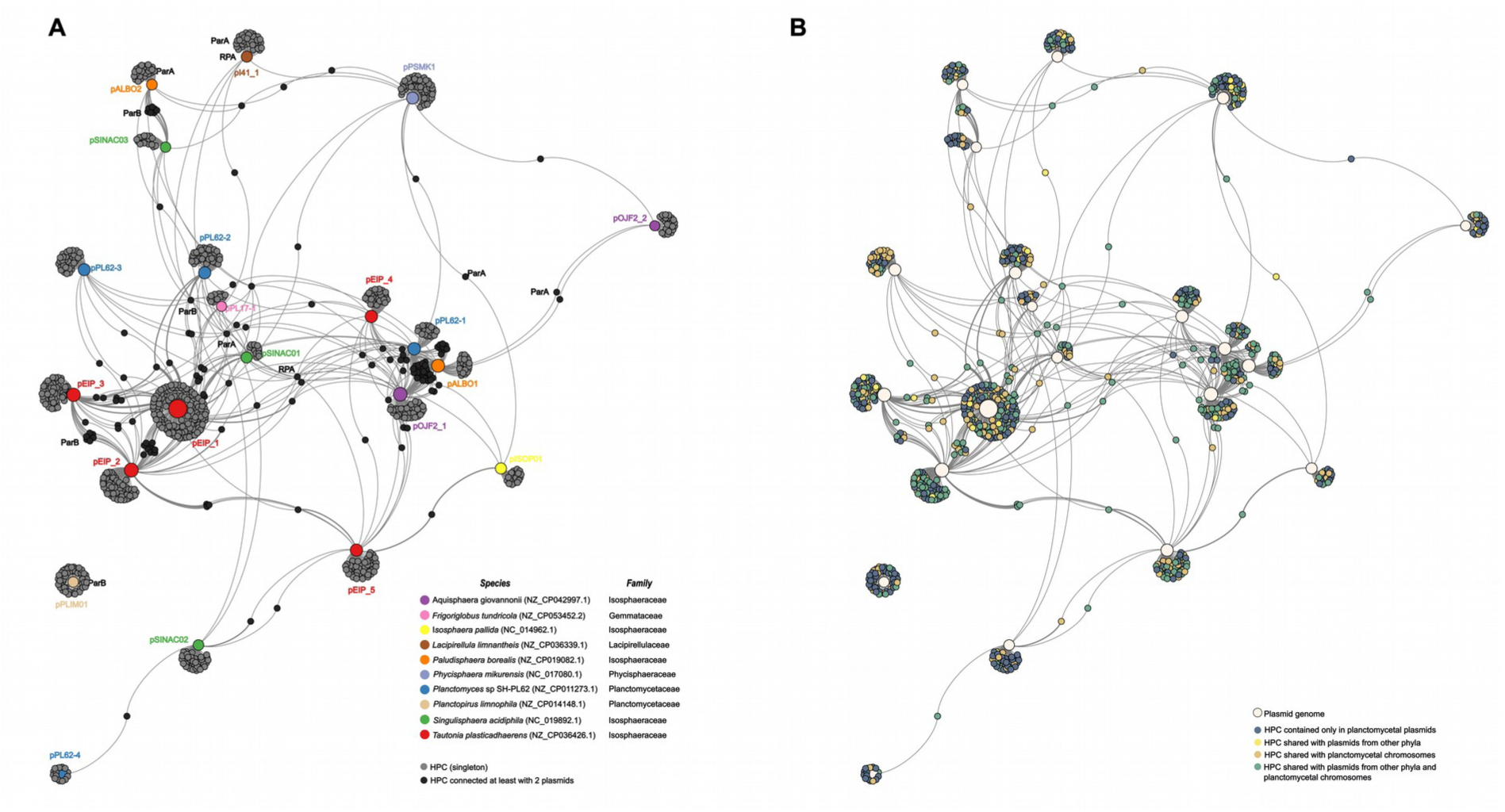
Proteome network of the Planctomycetes plasmidome. Bipartite network of homologous protein clusters at 30% identity and 60% coverage of planctomycetal plasmids. Nodes representing protein clusters are small, while nodes representing plasmids are larger and scaled according to plasmid size. **A)** Protein nodes are gray-colored, except for those containing members from at least two plasmids, which are depicted in black. Clusters containing RPA initiators and partition proteins (ParA, ParB) are indicated. Plasmids are colored by their host taxonomic family. **B)** The color of the protein nodes indicates if members of the HPC have homologs in the planctomycetal chromosome and/or in plasmids from other phyla.

Several HPCs emerged as central hubs within the network, connecting multiple plasmids (Figure 3A). Notable examples of these hub proteins included the replication initiation protein RPA (PF10134.12, 40% average identity, present in 12 plasmids), the partition Walker A motif-containing ATPase ParA (PF01656.26, with an average identity of 34%, associated with 13 plasmids), the centromere-like binding protein ParB (PF02195.21, with an average identity of 34%, present in 8 plasmids), and a tyrosine recombinase phage integrase (PF00589.25, with an average identity of 43%, observed in 9 plasmids).

Within the network, three plasmids originating from the *Isosphaeraceae* class (pPL62-1, pALBO1, and pOJF2_1) exhibited the most extensive interconnections, sharing a total of 37 HPCs. The shared HPCs encompassed a range of functions, including ABC transporters (PF00005.30, PF01061.27), as well as several CAZymes, such as glycosyl transferases (PF00534.23, PF06165.14, PF13439.9, PF13524.9, PF13579.9) and glycosyl hydrolases (PF17167.7). This observation aligns with the previously reported diverse repertoire of CAZymes associated with plasmids pPL62-1 and pALBO1 (Ivanova et al., 2017). Notably, pPL62-1 and pALBO1 have been previously documented to exhibit a high degree of similarity, sharing up to 51 homologous loci (Ivanova et al., 2017). This shared genetic content explains their substantial connectivity within our bipartite network, with 46 HPCs in common. In contrast, at the other end of the spectrum, plasmid pPL62-4 shared a single HPC, and pPLIM01, the only plasmid hosted out of the class *Planctomycetia*, remained entirely isolated within the network. Precisely in the most divergent planctomycetal plasmid, pLIM01 of *Planctopirus limnophila*, an essential gene, encoding a protein of unknown function (WP_013112526.1) was recently identified by transposon-directed insertion site sequencing (Rivas-Marin et al., 2023).

### Planctomycetal plasmids and chromosomes share a large proportion of biological functions

The majority of the plasmid proteins (823 out of 1,375) have a chromosomal homolog in Planctomycetes (Figure 3B and Supplementary Figure S2). They distributed in 581 HPCs when clustered at 30% identity and 60% coverage. The most abundant shared HPCs included glycosyl transferases (PF13692.9, PF00534.23, PF13641.9, PF13439.9, PF13524.9), epimerases (PF01370.24) and DDE transposases (PF13546.9, PF13586.9) (Supplementary Table S2). Thus, 552 plasmid proteins, distributed in 502 HPCs do not have a chromosomal homolog in Planctomycetes.

When comparing the Pfam content of planctomycetal plasmids and their host genomes, it was observed that 13 plasmids exhibited a substantial overlap, sharing at least 50% of the Pfam groups with the chromosome of their host (Supplementary Figure 1A). In contrast, 8 plasmids were characterized by a marked absence, ranging from 60% to 95%, of the Pfam content that was not present in the chromosome of their respective hosts. Among the protein families present in plasmids but absent in their hosts we found glycosyl transferases (PF03808.16, PF10111.12, PF13439.9, PF13506.9, PF13524.9, PF13579.9, PF13632.9, PF13641.9, PF13692.9; n=22 proteins), ParA partition protein (PF01656.26, PF13614.9; n=19 proteins), tetratricopeptide repeat (PF00515.31, PF07721.17, PF13374.9, PF13424.9, PF13428.9, PF13432.9; n=16 proteins), epimerases (PF00908.20, PF01370.24; n=14 proteins), homeodomain-like domains (PF13565.9; n=13 proteins), phage integrases (PF00589.25, PF02899.20; n=13 proteins), ParB partition protein (PF02195.21; n=12 proteins), response regulator receiver domain (PF00072.27; n=11 proteins), and DDE superfamily endonucleases (PF13358.9; n=8 proteins). Nevertheless, all these Pfam families were found chromosomally-encoded in other Planctomycetes.

### Shared functionalities found in plasmids within Planctomycetes and those originating from other bacterial phyla

Subsequently, we examined the evolutionary connections between plasmids originating from Planctomycetes and those from other phyla. First, to test whether the planctomycetal plasmids were similar to other bacterial plasmids, each plasmid was blastn-searched against the PLSDB database (version 2020_06_23), encompassing a total of 34,513 bacterial plasmids. Remarkably, no hits were retrieved with the sole exception being the 15 planctomycetal plasmids already included in PLSDB, which were identified when queried using the same plasmid sequences. Next, the proteome of planctomycetal and RefSeq212 plasmids was clustered at 30% identity and 60% coverage. This analysis unveiled 545 proteins derived from planctomycetal plasmids, distributed in 375 HPCs, that exhibited homology to counterparts in plasmids circulating out of Planctomycetes (Figure 3B, Supplementary Figure S3, and Supplementary Table S2). A majority of these HPCs, 82% of the clusters, contained planctomycetal proteins encoded exclusively in a single planctomycetal plasmid, in agreement with the low degree of connectivity between planctomycetal plasmids. Notably, a substantial majority of these shared proteins (535 out of 545) could be confidently classified into specific Pfam or COG groups (Figure 4, Supplementary Table S2). The most populated HPCs included homologs of the DDE superfamily endonuclease (PF13358.9), partition ParA protein (PF01656.26), phage integrases (PF00589.25), glycosyl transferases (PF00534.23), epimerases (PF01370.24), AAA+ ATPases (PF13614.9), homeodomain-containing (PF13565.9, PF13614.9), and RHS-containing (PF05593.17) proteins. These putative far-related homologs are found in a variety of plasmids from different PTUs, 97% of which exhibited a host range restricted to different taxonomic ranks into the *Enterobacterales* order (host range I-IV) (Supplementary Table S4). Noticeable, 42% of HPCs in common between planctomycetal plasmids and chromosomes lacked a homolog in plasmids from other phyla.

**Figure 4.**
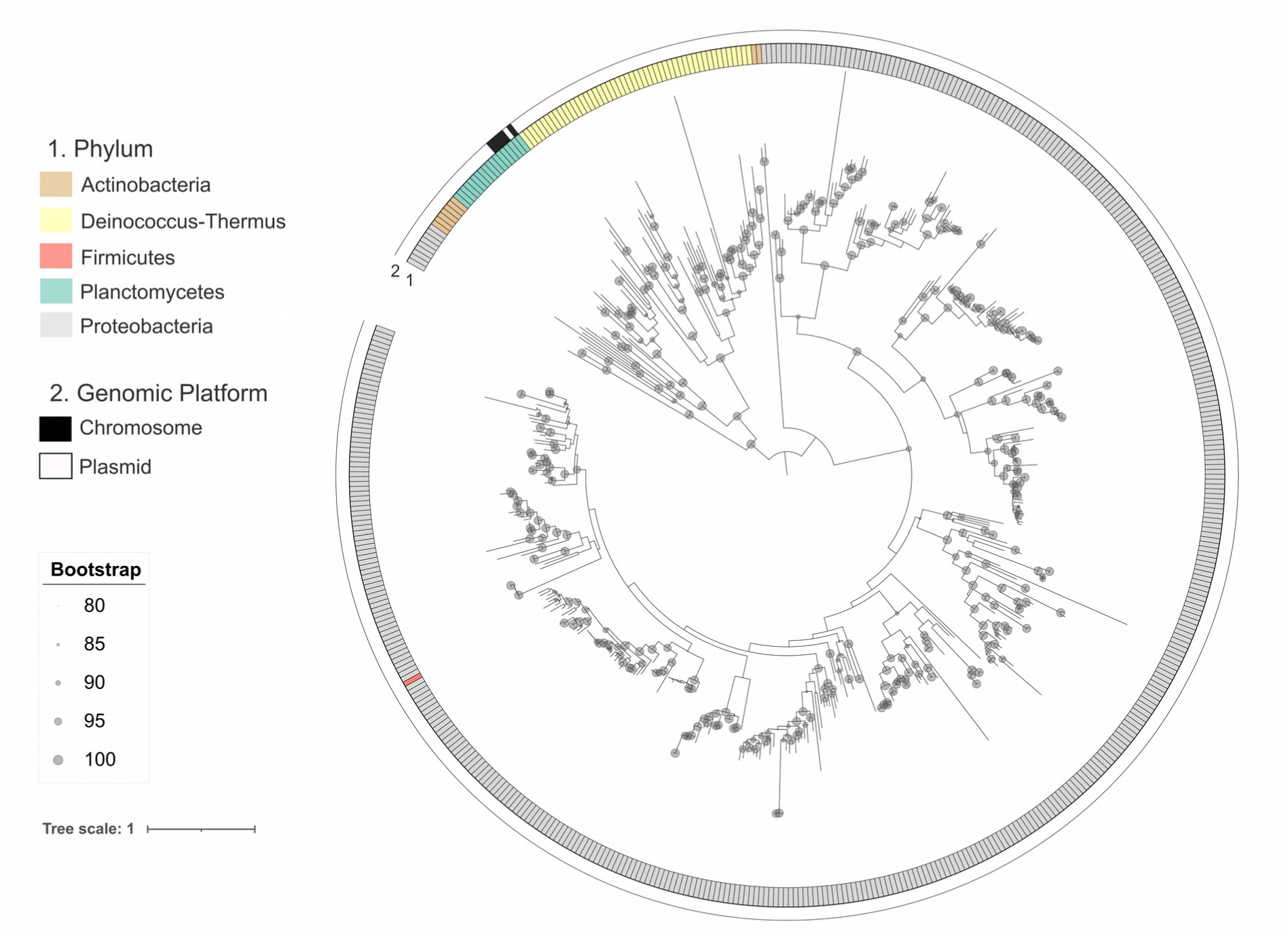
RPA protein phylogeny. A) Maximum likelihood tree based on 530 RPA amino acid sequences retrieved from plasmids in the RefSeq212 database, 13 from planctomycetal plasmids and 5 homologs detected in planctomycetes chromosomes, using the best-fit amino acid substitution model LG+F+R8. The tree was rooted at midpoint. Grey circles are placed on the branches supported by ultrafast bootstrap values >=80%. For each RPA homolog, genome and protein accession numbers are included at the tips. The different host phyla are shown in colors according to the legend (inner ring). RPA homologs detected in planctomycetes chromosomes are depicted in black in the outer ring. B) Close up of the planctomycetes clade of the RPA tree.

### Recent gene transfer events between plasmids and chromosomes in Planctomycetes

Instances of homology, particularly when subjected to rigorous criteria, serve as diagnostic indicators of recent genetic exchange events, inherently impeding the accumulation of substantial nucleotide alterations. In line with this approach, the proteomes from plasmids and chromosomes were subjected to a clustering process characterized by a 99% identity threshold and 100% coverage criterion. As a result of this analysis, we observed the presence of 12 proteins out of 1,375 encoded within planctomycetal plasmids that were concurrently identified in the chromosome of their host (Figure 7). Notably, these identified proteins primarily belonged to insertion sequences, including DDE transposases of the IS*701* (PF13546.9), IS*66* (PF03050.17), IS*4* (PF01609.24), and IS*4/5* (PF13340.9) families, as well as the IS*66* transposase accessory protein (PF05717.16), and three orphan ORFs. IS elements are recognized for facilitating the intracellular mobility of DNA, affecting not only their own genes but also those in their proximity (Partridge et al., 2018). Our results indicate the recent occurrence of transposition events between both genomic platforms in Planctomycetes.

### Diversity of planctomycetal plasmid modules

Plasmid backbones comprise distinct modules that encompass different functions in the biology of these mobile genetic elements, such as replication, stability, adaptation, and propagation. We searched for the presence of proteins encoded in these different modules in the Planctomycetes plasmid dataset.

### Plasmid replication

Plasmids can replicate autonomously because they contain an origin of replication (*oriV*) and generally encode the replication initiation protein (RIP) responsible for recognizing the cognate *oriV* and initiating plasmid replication (del Solar et al., 1998). An initial search of plasmid replicons by using prototypes from *Enterobacteriaceae* and Gram-positive bacteria as implemented in PlasmidFinder (Carattoli et al., 2014), or the origin of replication database implemented in DoriC 10.0 (Luo and Gao, 2019) did not render any hit in the planctomycetal plasmidome, suggesting that the replicons of these plasmids largely differ from those in other phyla, in agreement with the fact that attempts to introduce IncQ, IncP and pBBR replicons into Planctomycetes by conjugative matings failed to produce plasmid replicants (Jogler and Jogler, 2013). We then retrieved RIPs from the Planctomycetes plasmids by carrying a blastp search against known RIP families. For 14 out of the 21 planctomycetal plasmids a putative RIP was detected, 13 belonged to the RPA family (PF10134.12) and 3 to the HTH_36 family (PF00239.24) were found in three plasmids (Supplementary Table S1). These RPA homologs matched Pfam PF10134.12 with coverage values ranging from 58% to 93%. The average percentage identity of the 13 RPA proteins detected in Planctomycetes plasmids was 37.5%.

RPA is a single-stranded DNA binding protein that is required for multiple processes in eukaryotic DNA metabolism, including DNA replication, DNA repair, and recombination (Wold, 1997). It was functionally identified as a replication initiation protein in plasmid pBTK45 (GenBank Acc. no. EU585932.1) of the betaproteobacterium *Tetrathiobacter kashmirensis* strain WGT (Dam et al., 2009). We reconstructed the RPA gene phylogeny using the proteins retrieved with pfam PF10134.12 from the complete bacterial plasmid dataset in order to study the context of these planctomycetal RIPs regarding those from other phyla. A total of 530 RPA sequences were included in the phylogenetic tree (Figure 4). The RPA proteins detected in Planctomycetes clustered together in a separated clade from those present in the other phyla. Besides, RPA proteins from Planctomycetes did not clustered at 30% identity 60% coverage with the homologs out of this phylum. These facts suggest a large divergence of this subset of RPA replication proteins and its specialization in Planctomycetes, without recent interchanges with other hosts. The closest planctomycetal RPA homologs out of this phylum were found in Actinobacteria. Five RPA homologs were detected in the Planctomycetes chromosomes: two in *Lacipirellula parvula*, two in *Gemmata obscuriglobus*, and one in *Fuerstiella marisgermanici*. These chromosomal RPA homologs are located in a separate, more recent clade from most RPA encoded in plasmids, except for the RPA contained in plasmid pPL17-1, hosted in *Frigoriglobus tundricola*, which is intermingled with the chromosomal ones. It suggests that chromosomal RPA proteins are derived from plasmids that integrated into the chromosome.

Generally, the *oriV* sequences are located in close proximity to the replication-related genes, and are enriched in A+T content to facilitate the DNA unwinding (del Solar et al., 1998; Lilly and Camps, 2015). In the Planctomycetes plasmids, there is generally a decrease in the GC content in the region located immediately adjacent to genes encoding RPA initiators (500 bp upstream)(Supplementary Figure S5), suggesting a putative *oriV* sequence in that location. The *oriV* regions generally contain sequences repeated in tandem (iterons) that are recognized by the plasmid-encoded RIP and are critical to control the plasmid replication (del Solar et al., 1998). In the Planctomycetes plasmids where a member of the RPA replication protein family was identified, we performed a search for tandem repeats and found different kinds of iterons, either upstream, downstream or into the *rpa* gene (Supplementary Table S5).

### Plasmid stability

Segregational stability is essential for the plasmid maintenance in bacterial populations. Active partition is one of the plasmid segregation mechanisms encoded by the plasmids (Baxter and Funnell, 2014). Plasmid-encoded partitioning loci encode at least a *cis*-acting centromere-like site and two *trans*-acting proteins, a DNA-binding protein that binds specifically to the centromere-like site and an NTPase that forms filamentous structures (Ebersbach and Gerdes, 2005). The ATPase ParA (PF01656.26) and the DNA-binding protein ParB (PF02195.21) were detected in 19 and 12 plasmids, respectively. In 11 of these plasmids, the ParA and ParB homologs were detected in close genetic proximity, resembling a classical type I partition system (Supplementary Table S1, Figure 5). The average percentage identity of the detected ParA and ParB proteins was 29.5% and 34.4%, respectively.

**Figure 5.**
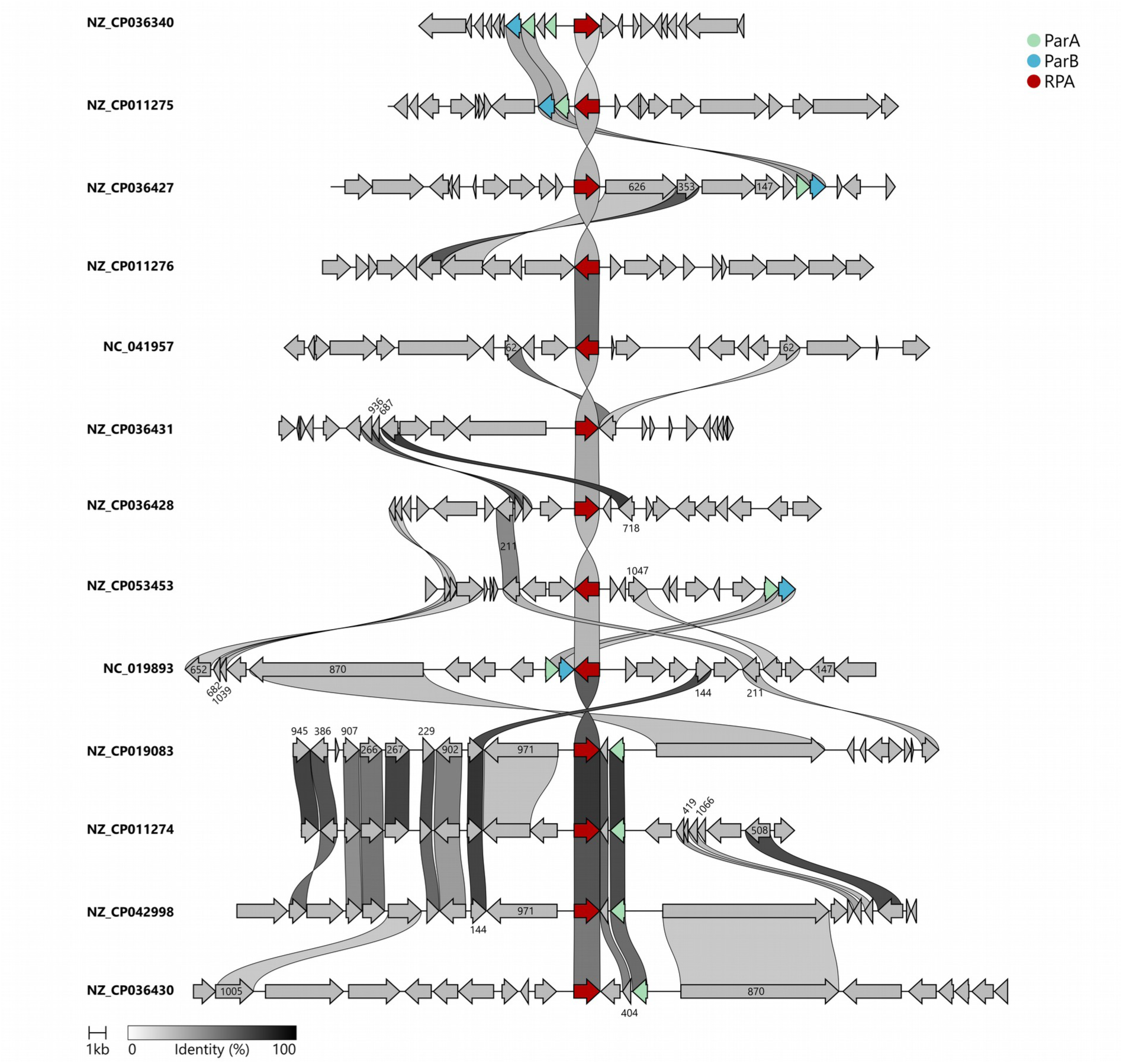
Genetic organization of the *rpa, parA* and *parB* homologs detected in planctomycetes plasmids. The genetic vicinity or the replication initiation gene *rpa* is compared between 13 planctomycetal plasmids by using clinker (Gilchrist and Chooi, 2021) with default parameters. Relevant genes are colored according to the legend.

Another strategy to maintain plasmid stability is post-segregational killing (Gerdes et al., 1986). This mechanism relies on toxin-antitoxin (TA) systems, which consist of a stable toxin that poisons the host cell and a much less stable cognate antitoxin that counteracts this detrimental effect, both components encoded by genes generally linked (Harms et al., 2018; Jurėnas et al., 2022). Plasmids encoding TA systems ensure their preservation in the bacterial population since cells lacking the plasmid, and thus the antidote, are eliminated (Díaz-Orejas et al., 2017). TA modules can be classified in eight different groups according to the molecular pattern of antitoxin and the mechanism of toxin neutralization (Jurėnas et al., 2022). The TADB 2.0 database (Xie et al., 2018) was used to retrieve putative TA modules in the Planctomycetes plasmid dataset. Among the different groups of TA, only type II TA proteins were detected. In type II TA systems, a direct protein-protein interaction between toxin and antitoxin blocks the action of the toxin (Gerdes et al., 1986). Putative type II TA modules were detected in 4 plasmids: AbiEii (PF08843.14) / AbiEi (PF17194.7+PF11459.11), RelE (PF06296.15) / MqsA (PF15731.8), HigB (PF09907.12) / HigA (PF01381.25), and Gp49 (PF05973.17) / HTH_37 (PF13744.9) (Table 1).

### Plasmid adaptation

Plasmid adaptative modules confer a beneficial advantage to the cells harboring them within a specific environment. Typically, these traits are associated with resistance to antimicrobials or metals, degradative pathways, and virulence factors. We screened the planctomycetal plasmids searching for resistance genes, and no hits meeting the threshold cutoff were identified. This contrasts with the fact that Planctomycetes are naturally resistant to many antibiotics (Cayrou et al., 2010; Godinho et al., 2019). Congruently, when searching in the Planctomycetes chromosomal dataset, 142 putative antimicrobial resistance traits were retrieved. Antibiotic efflux pumps were the most frequently detected, such as MexF (70), RosA (11), AbeS (11), KdpE (10), MexK (7) and MexQ (5) (Supplementary Table S6).

### Plasmid transfer

Plasmid transfer from a donor to a recipient cell via conjugation is another key feature of plasmid biology. Following introduction into the new host, the plasmid fate is determined by its replication capability, which may result in its retention as a self-replicating entity or its integration into the host chromosome, a state known as an Integrative and Conjugative Element (ICE) or Integrative and Mobilizable Element (IME) (Guglielmini et al., 2011). Plasmids and their integrated counterparts are categorized according to their potential for conjugation (Garcillán-Barcia and de la Cruz, 2013). They are classified as conjugative if they contain the entire conjugative apparatus, including relaxase and mating pair formation (MPF) systems. On the other hand, they are labeled as mobilizable if they only carry the coding information for a relaxase. Phylogenetically, relaxases are organized into nine distinct MOB families (Garcillán-Barcia et al., 2020), and the MPF systems are classified into eight different types (Guglielmini et al., 2014).

The notion that the presumed compartmentalization of Planctomycetes could act as a safeguard against horizontal gene transfer (HGT) has been considered (Pinos et al., 2016). However, this idea was dismissed, as no discernible differences were identified in terms of HGT quantity, proportion, or transfer partners when compared to other bacterial phyla (Pinos et al., 2016). Planctomycetes are not impermeable to HGT. Conjugative transfer to Planctomycetes from outside of this phylum has been documented. Plasmid pBF1, which was exogenously isolated from a marine microbial community (Partridge et al., 2018), was successfully transferred from *Pseudomonas putida* to *Planctomyces maris* (Dahlberg et al., 1998), and shuttle vectors were mobilized by the same plasmid pBF1 from *E. coli* to *Gemmata obscuriglobus*, *Blastopirellula marina, Gimesia maris (*formerly *Planctomyces maris*), and *Planctopirus limnophila*, where the transfer was detected by recombination (Rivas-Marín et al., 2016). Other attempts using the conjugative plasmid RP4 in matings from *E. coli* or *P. putida* to *Planctopirus limnophila* have failed (Fuerst, 2013).

To our knowledge, bacterial conjugation events among members of the Planctomycetes phylum have not been previously reported. A chromosomal relaxase gene (*traI*) was identified in a previous genomic analysis of *Gemmata obscuriglobus* (Jenkins et al., 2002). We identified a relaxase gene in four distinct plasmids, three of them relaxases falling within the MOB_F_ class and the remaining one classified as MOB_P_ (Supplementary Table S6). Out of these plasmids, only one, pPSMK1, hosted in *Phycisphaera mikurensis* (class *Phycisphaerae*), was found to encode a putative type T mating pair formation (MPF) system (MPF_T_), which is the most abundant type in phylum Proteobacteria (Guglielmini et al., 2014). The genetic arrangement of this putative conjugative system is illustrated in Figure 6. Notably, this putative conjugative system lacks the VirB1 and VirB7 components. VirB1 is a cell wall hydrolase that was found to be non-essential for T-DNA transfer (Berger and Christie, 1994; Fullner, 1998). VirB7 is a small fast-evolving lipoprotein (Guglielmini et al., 2014), which could explain that it went unnoticed.

**Figure 6.**
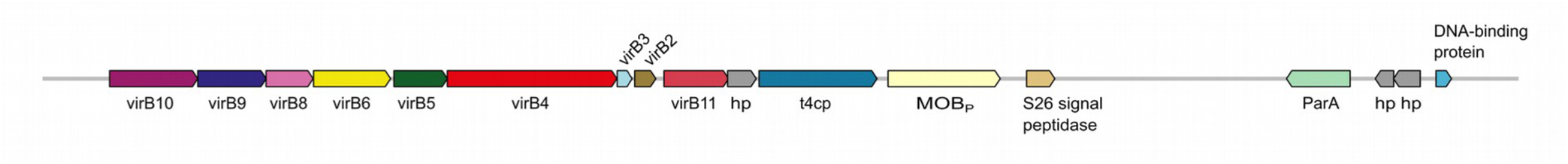
Genetic organization of the conjugative system present in plasmid pPSMK1. Genes encoded in the region comprising coordinates 60,558 to 80,714 of plasmid pPSMK1 (GenBank Acc. no. NC_017081.1) are shown.

Relaxases were also detected in Planctomycetes chromosomes (Supplementary Table S7), where two relaxases were found in the same genome in several cases: 29 MOB_F_ in 21 chromosomes and one MOB_P_ in one chromosome. A coupling protein (T4CP) was often detected along with the relaxase. No complete MPF system was detected in chromosomes, although some MPF proteins were found in 38 chromosomes.

## CONCLUSIONS

Planctomycetal plasmids seemed to have been largely coevolved with their hosts, as indicated by the high correlation between the GC content of the plasmids and their host genomes. 43% of the plasmid proteome could not be assigned either to a Pfam or COG category, highlighting the unexplored potential of distinct planctomycetal plasmids in offering proteins with diverse and novel functions. A large portion of the proteins encoded in the plactomycetal plasmidome (60%) showed to have a far-related chromosomal homolog. This and the fact that the functionally characterizable part of the planctomycetal plasmidome comprises a wide-range of bacterial gene functions suggest that most types of functions can be captured on plasmid DNA and become mobilized. Consistent with this, transposases from various IS elements have been identified to undergo recent mobilization events between planctomycetal chromosomes and their co-resident plasmids, providing a pathway for crosstalk between both genomic platforms.

Planctomycetal plasmids showed high diversity in gene content, a characteristic anticipated by the limited dataset of complete planctomycetal genomes. They were also very different from plasmids out of Planctomycetes, although remote homology could be detected in 40% of the planctomycetal proteome. In summary, 498 planctomycetal plasmid proteins, grouped in 451 HPCs, lacked a far-related homolog either in plasmids from other phyla or planctomycetal hosts. Thus, 36% of the proteins composing the planctomycetal plasmidome are exclusive.

We found that replication initiation proteins belonging to the RPA family are abundant in planctomycetal plasmids, and regions with characteristics compatible with origins of replication were identified in the proximity of these replication genes. Moreover, active type I partition systems were identified in several of these plasmids. These biological parts could be harnessed to design shuttle vectors able to be stably maintained in Planctomycetes. Finally, the very existence of mobilizable and conjugative elements in this phylum anticipates the possibility of bacterial conjugation among its members. To substantiate these findings, further experimental assays aimed at validating plasmid transfer via the putative planctomycetal conjugative system could offer valuable insights into the dynamics of bacterial conjugation within this phylum.

## METHODS

### Planctomycetes dataset

The whole set of complete genome sequences of *Planctomycetota* available at NCBI (June 2022), 83 full genomes (83 chromosomes and 21 plasmids), was retrieved (Supplementary Table S8). Bacterial plasmids (43,874 sequences) were obtained from the NCBI RefSeq212 database. To remove sequences incorrectly tagged as plasmids, plasmid records were filtered by their description, using the regular expression “*contig|\sgene(?!tic| ral|rat|ric)|integron|transposon|scaffold|insertion sequence|insertion element|phage|operon| partial sequence|partial plasmid|region|fragment|locus|complete (?!sequence|genome| plasmid|\.|,)|(?<!complete sequence,) whole genome shotgun|artificial|synthetic|vector*”. The curated plasmid dataset comprised 43,217 sequences.

### Genome phylogenetic analysis

PhyloPhlAn v3.0.67 (Asnicar et al., 2020) was used for detecting the 400 most universal prokaryote markers in the set of 83 planctomycetal chromosomes and producing a concatenated multiple sequence protein alignment, using the parameters *-d phylophlan –– accurate ––diversity high*. The resulting alignment was in turn trimmed with Trimal v1.2 (Capella-Gutiérrez et al., 2009), using option –*automated1*. A total of 8,253 aligned amino acid positions were used to build a Maximum-Likelihood (ML) tree with IQ-TREE v1.6.12 (Nguyen et al., 2015) with the best-fit amino acid substitution model LG+F+R5 (Kalyaanamoorthy et al., 2017) and 1,000 ultrafast bootstrap replicates (Hoang et al., 2018). *Chlamydia pecorum* E58 chromosome (GenBank Acc. No. NC_015408.1) was included in the analysis as an outgroup. The phylogenetic tree was visualized with the iTOL web platform (https://itol.embl.de/) (Letunic and Bork, 2019).

### Protein phylogenetic analysis

For the RPA replication initiation protein phylogeny, amino acid sequences were aligned using MAFFT v7.271 (Katoh et al., 2002) and the alignment was trimmed with Trimal v1.2 (*-automated1*). IQ-TREE tool was also used to generate a maximum likelihood (ML) tree with 1,000 ultrafast bootstrap replicates and the best-fit amino acid substitution model LG+F+R8. The phylogenetic tree was rooted at midpoint and visualized with iTOL.

### Comparison of GC content

The GC content of plasmids and chromosomes was calculated with home-made scripts in Python v3.7.1. GC correlation between plasmids and chromosomes was calculated with the scipy.stats.pearsonr function from Python. Plots and graphs were generated with the R package ggplot2 v3.3.6. For GC content analysis of the adjacent region of replication initiation protein genes, the Kruskal-Wallis test was applied, using the Python base function scipy.stats.kruskal.

### Characterization of the planctomycetal plasmid proteome

Plasmid and chromosomal proteomes of the *Planctomycetota* phylum were mapped to Pfam (Mistry et al., 2021) and eggNOG v5.0 (Huerta-Cepas et al., 2019) databases. Hidden Markov Models (HMM) for 19,632 protein families contained in Pfam-A 35.0 database released in November 2021 were downloaded (http://pfam.xfam.org/) and used to search for protein families in our dataset with the *hmmscan* function of HMMER suite v3.1b2 (Eddy, 2011) (parameters *-E 0.0001 ––domE 0.0001 ––incE 0.0001 ––incdomE 0.0001*). Only hits covering at least 80% of the protein profile were recorded in Supplementary Table S2. eggNOG-mapper v2.1.9 (http://eggnog-mapper.embl.de/) was used to identify putative Cluster of Orthologous Groups (COGs) with the bacteria optimized database (e-value <= 10^-4^ and 80% subject coverage). To search for plasmid replicons we used several tools: PlasmidFinder (Carattoli et al., 2014), DoriC 10.0 (Luo and Gao, 2019), and blastp (e-value <=10^-5^) against the replication initiation protein database implemented in PLACNETw (Vielva et al., 2017)(Supplementary Table S9). A blastp (e-value <=10^-5^) search was also used to detect putative toxin-antitoxin (TA) systems for the 6 subtypes available in the TADB 2.0 database (Xie et al., 2018). Antibiotic resistance genes were screened based on CARD (McArthur et al., 2013) through a blastp search (50% amino acid identity, 60% sequence coverage, e-value <=10^-5^). MOBScan (Garcillán-Barcia et al., 2020) was used to search for the presence of relaxases both in plasmid and chromosomal genomes, while MacSyFinder (Abby et al., 2014) was employed to detect putative mating pair formation homologs.

### Clusterization methods

Average Nucleotide Identity with a 50% length threshold (ANI_L50_) was used to determine the similarity of the nucleotide sequences of the Planctomycetes plasmids with a 50% threshold in the length of the smaller genome in the pair (Redondo-Salvo et al., 2020). In that way, only plasmid pairs that exhibited >=70% nucleotide identity in at least 50% of the smaller genome were considered for scoring ANI_L50_, while those not fulfilling the threshold were assigned ANI_L50_=0. Blastn searches using planctomycetal plasmids as queries against the PLSDB database (version 2020_06_23) were used to detect similar plasmids in other phyla (e-value 0.001, 50% nucleotide identity, and 70% sequence coverage). Homologous Protein Clusters (HPCs) were generated with MMseqs2 (Steinegger and Söding, 2017) at 30% identity and 60% alignment coverage or 99% identity and 100% coverage (parameters *--cov-mode 0 –cluster-mode 0 ––cluster-reassign*). Bipartite networks containing plasmid and protein cluster nodes were constructed. Networks were visualized with Gephi v0.9 (https://gephi.org/).

## Acknowledgments

This work was supported by the Spanish Ministry of Science and Innovation (Grant MCIN/AEI/10.13039/501100011033 PID2020-117923GB-I00 to MPG-B), the Spanish Ministry of Universities (predoctoral contract FPU18/02647 to MMQC), and the “Moore-Simons Project on the Origin of the Eukaryotic Cell” (Grant No. 9733/ DOI https://doi.org/10.37807/GBMF9733 to DPD).

