## Supplementary figures for "Specificities and commonalities of the Planctomycetes plasmidome"

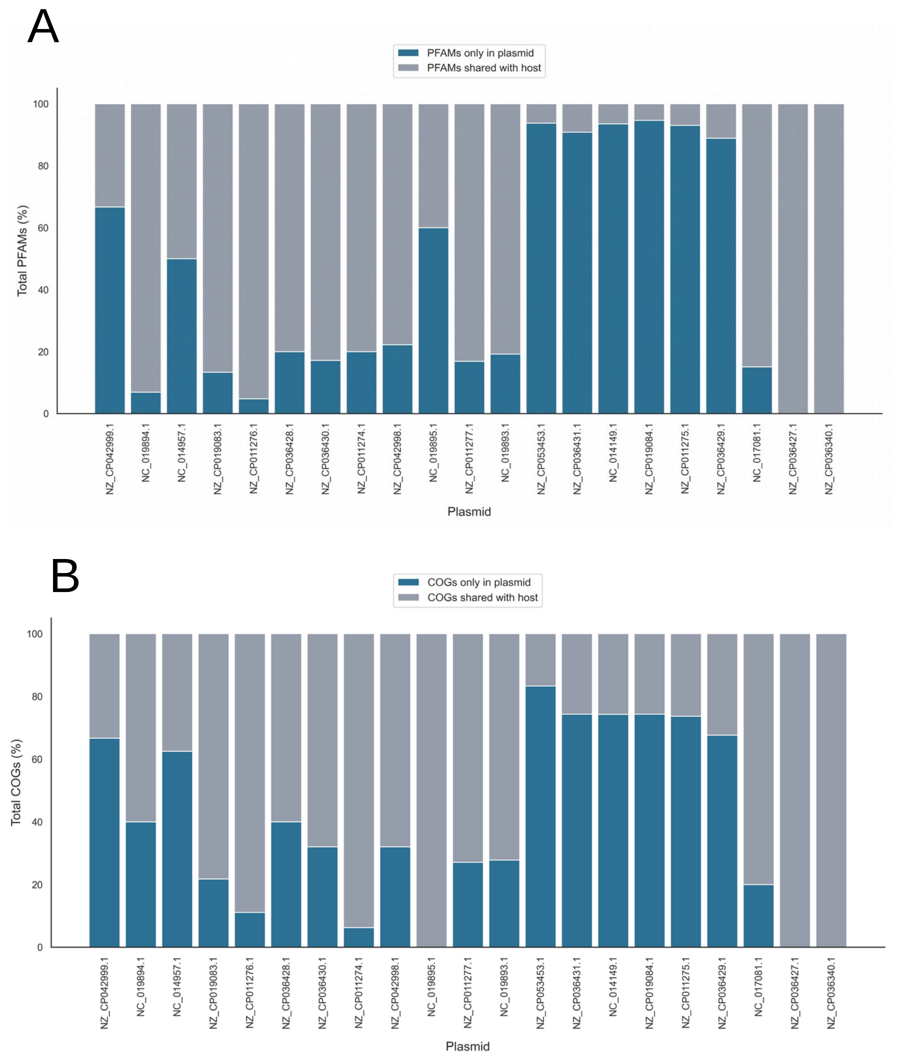


**Supplementary Figure S1: Functional annotation of the planctomycetal plasmid proteome.** The percentage of proteins families of Pfam database (A) and cluster of orthologous groups (COG) of eggNOG database (B) found for each plasmid in the Planctomycetes phylum.

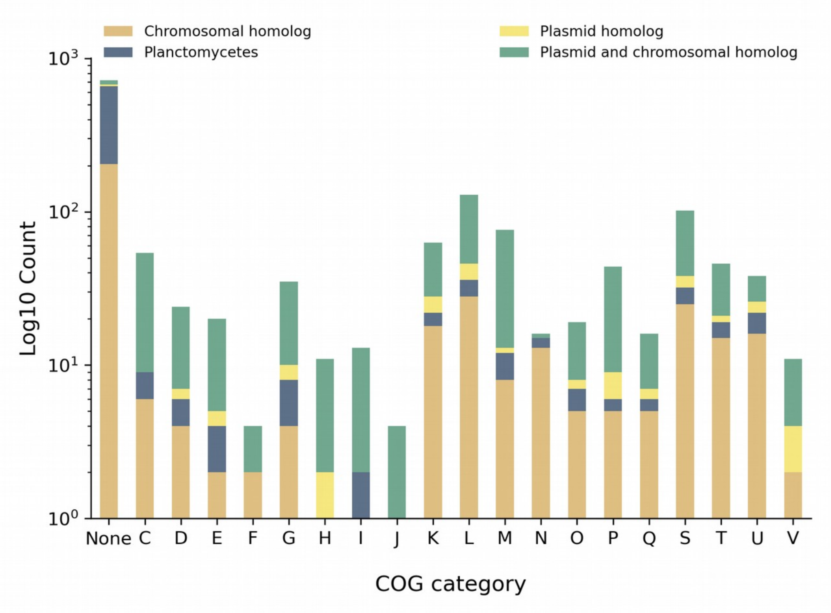


**Supplementary Figure S2: Specific and shared proteins of the planctomycetal plasmid proteome.** For each COG category, the abundance of plasmid proteins is indicated. Each bar is subdivided in different colors: homologs in the planctomycetal chromosome (orange), homologs in plasmids from other phyla (yellow), homologs in both planctomycetal chromosome and plasmids from other phyla (green), planctomycetal plasmid proteins without homologs (dark blue).

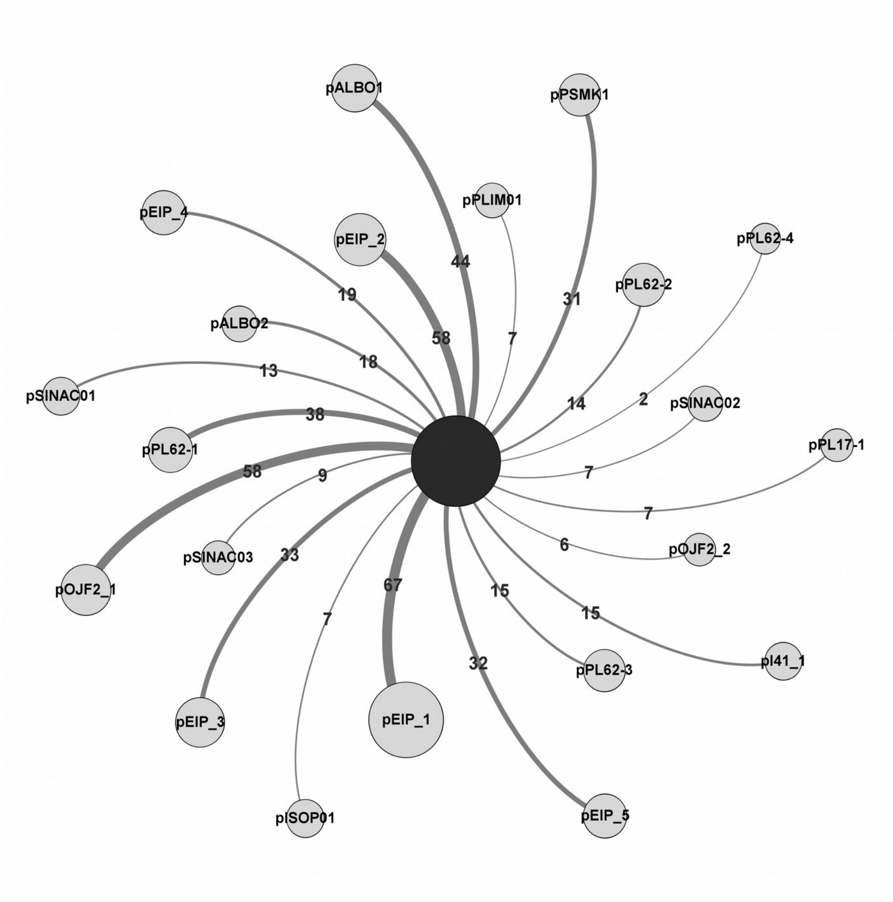


**Supplementary Figure S3: Proteins shared between planctomycetal plasmids and those from other phyla.** For each planctomycetal plasmid the number of proteins that clustered with proteins from plasmids out of Planctomycetes is shown.

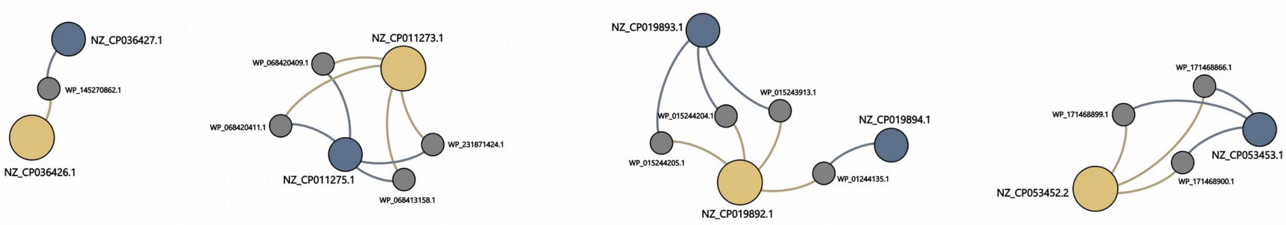


**Supplementary Figure S4: Network of recent HGT events between planctomycetal plasmids and their host genomes.** Twelve HPCs (colored in grey) are connected to cohabiting plasmids (blue) and chromosomes (yellow) at 99% identity and 100% coverage. The accession numbers of plasmids, chromosomes, and a representative protein of each HPC are indicated.

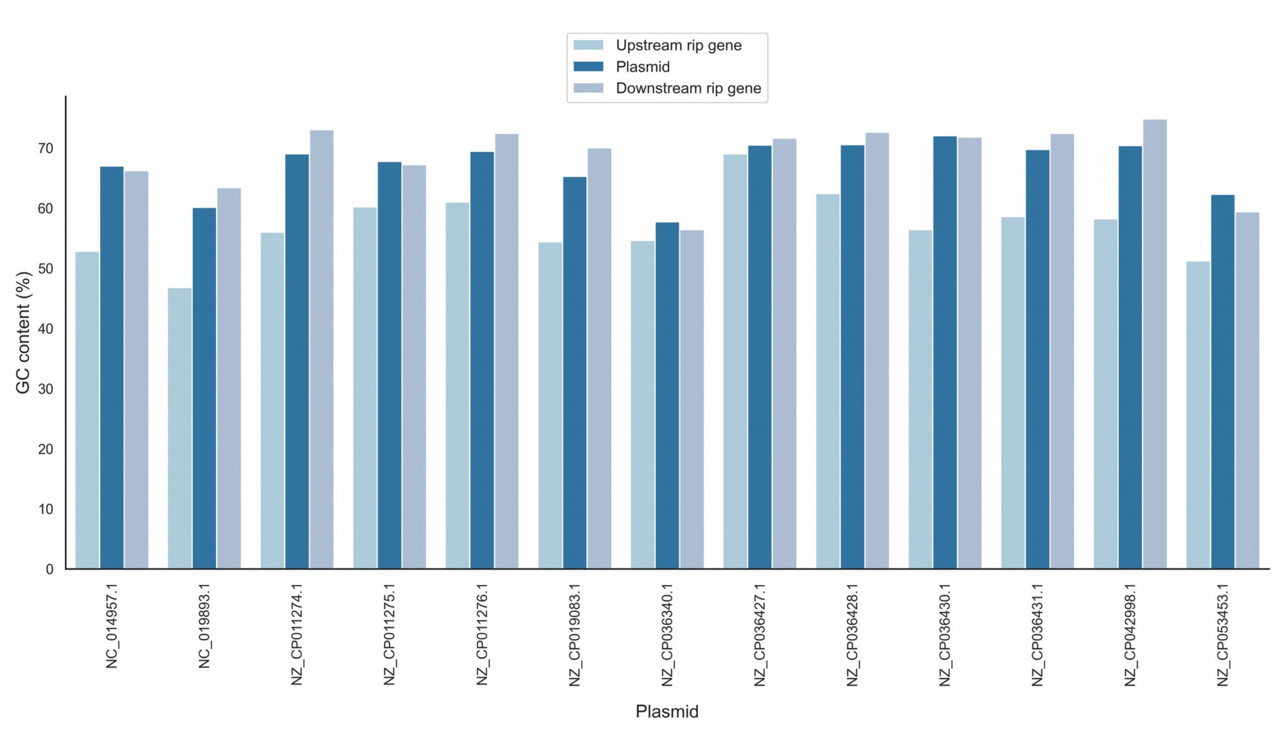


**Supplementary Figure S5: GC content in the neighborhood of the *rpa* gene.** For each plasmid encoding an RPA homolog, the GC content of the upstream and downstream region (500bp) of the *rpa* gene are compared with that of the chromosome.
